# REV-ERB Agonism Improves Liver Pathology in a Mouse Model of NASH

**DOI:** 10.1101/2020.06.29.177378

**Authors:** Kristine Griffett, Gonzalo Bedia-Diaz, Bahaa Elgendy, Thomas P. Burris

## Abstract

Non-alcoholic fatty liver disease (NAFLD) affects a significant number of people worldwide and currently there are no pharmacological treatments. NAFLD often presents with obesity, insulin resistance, and in some cases cardiovascular diseases. There is a clear need for treatment options to alleviate this disease since it often progresses to much more the much more severe non-alcoholic steatohepatitis (NASH). The REV-ERB nuclear receptor is a transcriptional repressor that regulates physiological processes involved in the development of NAFLD including lipogenesis and inflammation. We hypothesized that pharmacologically activating REV-ERB would suppress the progression of fatty liver in a mouse model of NASH. Using REV-ERB agonist SR9009 in a mouse NASH model, we demonstrate the beneficial effects of REV-ERB activation that led to an overall improvement of hepatic health by suppressing hepatic fibrosis and inflammatory response.

## Introduction

Among the metabolic disorders, non-alcoholic fatty liver disease (NAFLD) is considered a hepatic manifestation of metabolic syndrome (MetS) and it is one of the prominent health challenges of the twenty-first century as NAFLD is the most prevalent liver disease worldwide affecting 25-30% of the general population and its prevalence could reach 70-90% in specific populations with comorbidities such as morbid obesity or type 2 diabetes mellitus [1–2]. NAFLD can often progress to non-alcoholic steatohepatitis (NASH), which is associated with progressive liver disease [1]. NASH has been mainly associated with higher morbidity and mortality than other diseases in the NAFLD spectrum and, although there are pharmacological therapies under clinical investigation for treatment of NASH [2], no drugs are approved by the Federal Drug Administration (FDA) or the European Medicines Agency (EMA) for the NASH treatment [3].

Nuclear receptors (NRs) are transcription factors generally activated by ligands and involved in diverse biological processes such as cell growth and differentiation, apoptosis, gene expression during tumor formation and metabolism. They bind to specific sequences of DNA allowing them to regulate the expression of adjacent genes. Many diseases including NASH are directly or indirectly related to nuclear receptor signaling and many NRs have become favored targets for drug discovery [4]. NRs play an important role in liver diseases and they are key modulators in the onset and progression NAFLD, including the peroxisome proliferator-activated receptors (PPAR) α/β/γ; liver X receptors (LXR) α/β; farnesoid X receptors (FXR); constitutive androstane receptor (CAR); and pregnane X receptor (PXR). All of these NRs form obligate heterodimers with retinoid X receptor (RXR) α/β/γ in order to modulate corresponding target genes in the nucleus [5,6].

REV-ERB nuclear receptors (REV-ERBα and REV-ERBβ) are transcriptional repressors that regulate a variety of physiological processes including lipogenesis, inflammation, circadian regulation, and muscle regeneration and are expressed in all tissues but has significantly higher expression in liver, skeletal muscle, adipose tissue, and brain [7]. Although REV-ERBs play a regulatory role in hepatic metabolism, inflammation and lipogenesis, these NRs have yet to be validated as a potential therapeutic target for liver disease [8–11]. Here, we show that REV-ERB pan-agonist SR9009 treatment in *ob/ob* mice fed a high-fat, high-fructose (NASH) diet has beneficial effects and may provide some translational groundwork for further developing REV-ERB agonists for metabolic diseases, specifically NAFLD.

## Materials and Methods

### Animals

Animal studies were performed as previously described [12–14]. Briefly, six-week old B6 V-Lep^ob^/J (*ob/ob*) male mice were purchased from Jackson Labs (Bar Harbor, ME). Upon receipt, mice were housed individually in standard cages with huts and immediately placed on NASH diet (D09100301; Research Diets) [15]. Mice were maintained on this diet throughout the experiment. Mice were handled and weighed weekly while acclimating to the diet. At 12-weeks of age, mice were assigned into weight-matched groups (n = 7) and dosing began. Mice were weighed daily and food-intake was monitored daily. At the termination of the study, mice were fasted and euthanized by CO_2_ and blood was collected by cardiac puncture for clinical chemistry analysis at Scripps Florida Metabolism Core (Roche COBAS instrument) and ELISA analysis (EMD Millipore). Tissues were collected and flash-frozen in liquid nitrogen for gene expression, or placed in 4% Paraformaldehyde (PFA) in PBS for paraffin-embedding (Saint Louis University Histology Core) or 10% Neutral-Buffered Formalin (NBF) for cryo-sectioning. For intraperitoneal glucose tolerance test (ipGTT), mice were transferred to clean cages and fasted for 12 hours overnight. Mice were weighed, baseline blood glucose was taken, then mice were given an injection of glucose solution in PBS (2g/kg body weight). Blood glucose levels were subsequently repeated at 30-, 60-, 90-, and 120-minutes following the injection of glucose. Mean blood glucose levels (mg/dL) are reported as well as the area under the curve (AUC) which was analyzed by two-tailed student’s t-test in Graphpad prism. Following the ipGTT, mice were given access to food *ad libitum*. All animal work was approved by the Institutional Animal Care and Use Committee (IACUC) at Washington University in St. Louis (Protocol #20180062).

### Compounds and Dosing

SR9009 was formulated as 100mg/kg at 10mg/ml in 5% DMSO, 15% Cremophore EL (Sigma), 80% PBS as previously described [16]. Both vehicle (5% DMSO, 15% CremophoreEL (Sigma), 80% PBS) and SR9009 were filter sterilized (Millipore Steriflip) prior to dosing. Mice were given once daily i.p. injections within an hour of “lights on” (ZT0-ZT1). Dosing was performed for 30 days by the same researcher.

### Gene Expression Analysis

Total RNA was isolated from liver using the trizol (Invitrogen) method [12]. Samples were analyzed by QPCR using Fatty Liver and Fibrosis QPCR array plates (Bio-Rad; 384-well format) and Bio-Rad supplied SYBR reagents (per manufacturer’s protocol). Each sample was run in duplicate and analyzed on the PrimePCR software supplied by Bio-Rad. Multiple reference genes were utilized (including Gapdh, ActinB, and Cyclophillin) for analysis [14]. Results were plotted in GraphPad prism software as Gene Regulation using mean +/-SEM.

### Histology and Pathology Analysis

Livers were placed in 4% PFA at 4°C overnight and then were paraffin-embedded and sectioned at 10 μm onto slides at the Saint Louis University Histology Core Facility. H&E and Masson’s Trichrome staining were also performed at the core as a fee-for-service [12,14,17].. Stained sections (both H&E and Masson’s Trichrome) were sent to Reveal Biosciences (San Diego, CA) for quantification of fibrosis, inflammation, steatosis, hepatocellular ballooning, and presence of Mallory Bodies utilizing AI-based digital pathology and image processing. Briefly, whole slide images were generated using 3D Histech Pannoramic SCAN and uploaded into imageDx™ software. Each scanned image was assessed for quality and accuracy, then machine learning algorithms were applied to perform automated image quality control assessments and quantitative measurements of disease features across the entire tissue. For steatosis analysis, lipid regions within the H&E slides were identified and quantitated as a percent of the total image analysis area. For hepatocyte ballooning, these hepatocytes were identified and quantitated as the number of ballooning cells within the total image analysis area of the H&E stained sections. Positive ballooning was identified based on cell diameter and prescence of disrupted cytoskeletal structure. The number of ballooning cells was counted and expressed as density across the total area. Inflammatory cells were identified and quantitated on H&E slides as the number of immune cells within the total image analysis area. The number of immune cells was counted and measured as the immune cell density, total number, and area of immune cells. For fibrosis, collagen and extracellular matrix fibers were identified in the Massson’s Trichrome-stained slides and quantitated as a percent of the total image analysis area. H&E sections were also used to identify the presence of Mallory bodies within the total image analysis areas.

### Statistics

All data are expressed as mean +/-SEM (n = 4 or greater). All expression statistical analysis was performed using ANOVA with Tukey’s post-hoc analysis in GraphPad prism software. Weekly mouse weights and food intake data was analyzed by two-way ANOVA followed by Sidak’s multiple comparisons at the 95% confidence level. For ipGTT, area under the curve (AUC) was generated and the data was analyzed by two-tailed student’s t-test in GraphPad prism software. Sample data generated by Reveal Biosciences was entered into Graphpad prism software and analyzed for statistical significance using an unpaired student’s t-test (2-tailed). P-values are reported as follows: * p ≤ 0.05, ** p ≤ 0.01, *** p ≤ 0.001, and **** p ≤ 0.0001.

## Results

Given that REV-ERBs have been demonstrated to play a regulatory role in hepatic lipid metabolism [18] as well as inflammation [16,18,19], we sought to examine the effects of pharmacologically activating REV-ERB in a mouse model of NASH and determine whether REV-ERB may be a therapeutically relevant target. We opted to utilize a diet-induced NASH model using *ob/ob* mice as previously described [14,15] as the time period for development of NAFLD with fibrosis (NASH) is relatively short as compared to other models. During the acclimation and NASH development period, mice were fed a diet that contains Primex as a trans-fat source, fructose, and cholesterol *ad libitum* and monitored for weight gain and food intake. These parameters were also monitored daily throughout the dosing period to validate that any weight-loss was not due to loss of appetite. As shown in Figure 1A, both groups (Vehicle and SR9009) gained weight throughout the experimental period, however the SR9009-treated group gained weight at a consistently slower rate. The slower weight gain was not due to lower food intake in the SR9009-treated mice since this group consistently consumed the same amount of food as the vehicle group (Figure 1B). After 30-days of dosing was completed, mice were euthanized, and we performed a variety of tissue and plasma analyses to determine whether SR9009 treatment had a beneficial effect in this model. While we did not see a significant effect in liver weight as a percentage of total body weight (S1 Fig), we did observe a decrease in the fat mass of the SR9009-treated animals, while lean mass was unchanged (Fig 1C). We were interested in determining whether SR9009 had any utility in improving the hepatic health of these mice. In order to make this determination, we performed clinical chemistry analysis on blood plasma samples to examine liver enzyme levels. ALT was significantly decreased in the SR9009-treated group (Fig 1D). While not statistically significant, AST levels in the SR9009 group were also trending lower than the vehicle group consistent with a benefit due to SR9009 treatment (S1 Fig). These data suggest that that the amount of liver damage due to the diet may be suppressed by treatment with REV-ERB agonist SR9009. In addition to effects on hepatic enzymes, we also observed a significant decrease in fasted blood-glucose levels in the SR9009-treated mice (Fig 1E). The *ob/ob* mouse model is typically hyperglycemic and addition of the high fat/high fructose diet hyperglycemia can be particularly prominent. During the third week of dosing, we performed an ipGTT on the mice (Fig 1F) and observed that while both groups of mice were hyperglycemic, the SR9009-treated group responded better to the bolus of glucose. In fact, the difference in AUC was statistically significant between the two groups (Fig 1F right panel) as analyzed by a two-tailed student’s t-test (*p = 0*.*027*), although final plasma insulin levels were not affected by the treatment (S1 Fig). Thus, our observation that of lowered hyperglycemia was promising and potentially relevant to human NASH patients who often present with co-morbidities such as obesity and diabetes [20–24].

**Figure 1:**
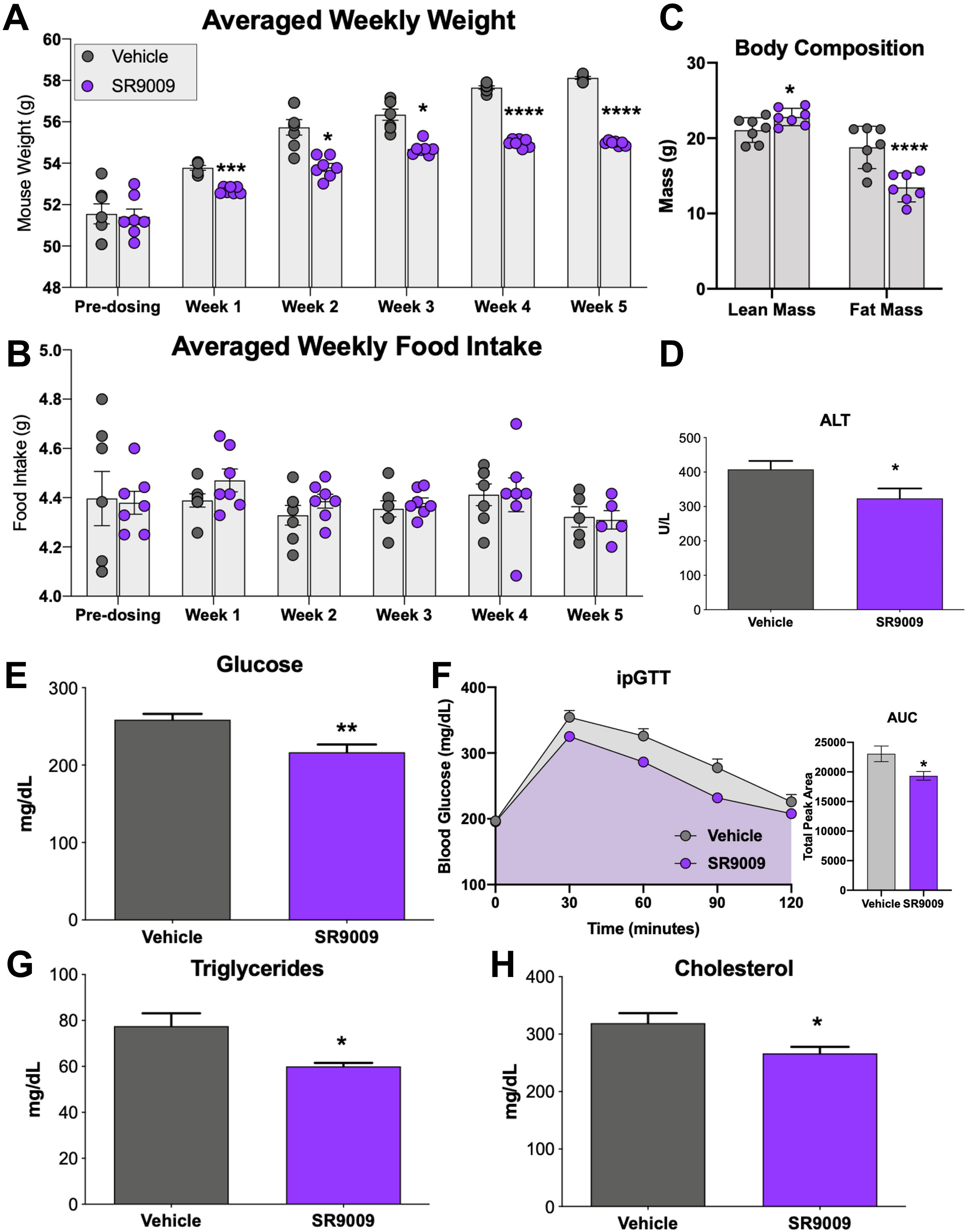
REV-ERB agonist treatment of ob/ob NASH diet-fed mice. Mouse weight (A) and food intake (B) were recorded daily and averaged weekly for each group. At the termination of the experiment, mice were euthanized and blood and tissues were collected for analysis. Panel C shows body composition of lean and fat mass for each group. ALT (D), fasting blood glucose (E), ipGTT and area under curve (AUC) (F), circulating triglycerides (G), and circulating cholesterol levels (H) were also analyzed from blood plasma.

Hyperlipidemia is also a comorbidity in NASH patients, therefore we also analyzed circulating triglyceride levels (Fig 1G) and total cholesterol levels (Fig 1H) in both mouse groups. We also observed significantly reduced total protein in the plasma of SR9009 mice as compared to vehicle-treated mice (S1 Fig). Our overall impression from the clinical chemistry data and mouse observations is that SR9009 treatment may have beneficial effects in a NASH model.

As the SR9009 group maintained a lower body weight throughout the dosing period and had significantly decreased circulating lipid levels in blood plasma, we investigated whether the SR9009-treated mice had reduced hepatosteatosis (both macrovesicular and microvesicular) [12,14,17]. Figure 2A and 2B show the total lipid accumulation and percentage of tissue covered in lipid accumulation based on the H&E stained liver sections for both vehicle- and SR9009-treated animals. There is no change in either macrovesicular or microvesicular steatosis (S1 Table), suggesting that the beneficial effects observed in this mouse model may not be attributable to REV-ERB’s role in reduced fat mass.

**Figure 2:**
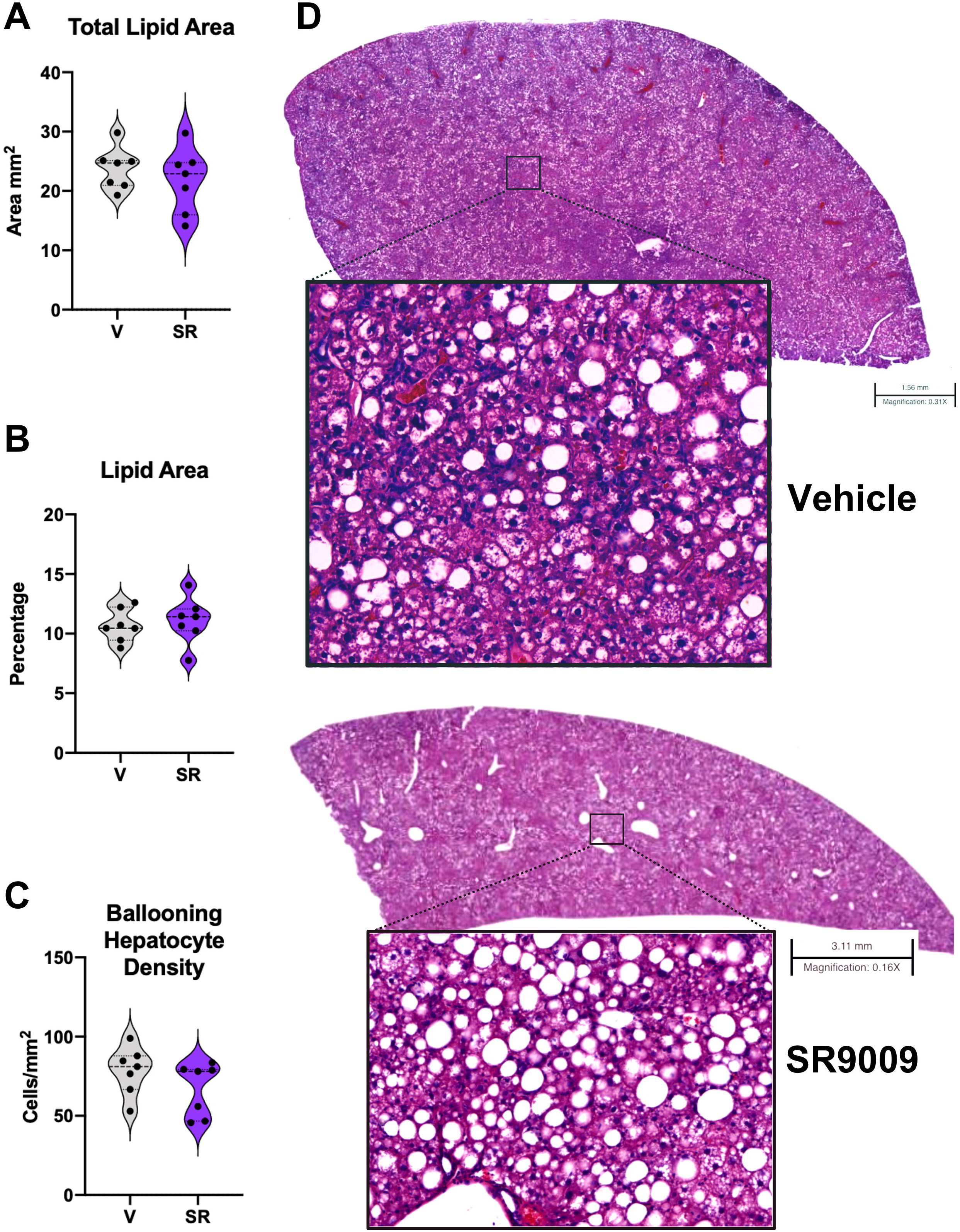
Hepatosteatosis in *ob/ob* mice maintained on a NASH diet does not appear affected in SR9009-treated group. Digital pathology analysis suggests that SR9009 treatment in NASH mice may not suppress total hepatic lipid area (A) or total percentage of lipids within the entire section (B). While steatosis appears unaffected by SR9009 treatment, the number of ballooning hepatocytes is decreased in the treated mice although the effect does not reach significance (C). (D) H&E sections of representative livers for vehicle (top) and SR9009 (bottom). Scale bar for vehicle section indicates 1.56 mm. Scale bar for SR9009 section indicates 3.11 mm. For steatosis quantitation, macro droplet > 65 μm^2^; micro droplet ≤ 65 μm^2^.

To validate the clinical chemistry data suggesting that the SR9009 reduced liver damage and improved liver health, quantitated hepatocyte ballooning density from H&E stained liver sections. As shown by Fig 2C, SR9009-treated mice have a lowered density of hepatocyte ballooning as compared to vehicle-treated mice. While this was not statistically significant (*p = 0*.*0903*), it suggests that treatment with SR9009 may have slowed the progression of NASH in these animals. We were intrigued that REV-ERB agonism in this model did not significantly affect steatosis in the liver with its known role in lipid metabolism, but appeared to suppress circulating cholesterol and reduce overall body fat mass. Therefore we assessed the expression levels of several known REV-ERB target genes involved in hyperlipidemia (*Dhcr24, ApoE*, and *ApoC3*) by QPCR and observed a significant decrease in these genes (S2 Fig) suggesting that SR9009 activation of REV-ERB is suppressing cholesterol and lipid metabolism but may not have efficacy in this model in which treatment started well after the development of NASH began.

While hepatic steatosis was not significantly affected by SR9009 treatment in this model, we further investigated clinical chemistry findings that SR9009 improves hepatic health by quantitating fibrosis and immune cell infiltration in liver sections (Fig 3A). We first evaluated the Masson’s trichrome stained sections and identified the collagen and extracellular matrix fibers stained with deep blue within the whole slide image. Positive collagen fibers were then visualized using an overlaid green mask and measured as a percentage of the total image analysis area (S3 Fig). The calculated percent area of positive collagen (fibrosis) was significantly reduced in the SR9009-treated mice (Fig 3B) suggesting that the beneficial hepatic effects from REV-ERB agonism in this model may be due to suppressed fibrosis. Further analysis of H&E staining also indicated a reduced number of inflammatory foci and immune cell count, although these did not reach significance in our analysis (Fig 3C-D). Additionally, H&E sections indicated that all liver samples in this study showed the presence of Mallory Bodies, however the overall morphology of the SR9009-treated livers appeared improved as compared to the vehicle group (S1 Table). Overall, these results suggest that SR9009 treatment halted the progression of NASH in this mouse model by reducing hepatic inflammation and suppressing fibrosis.

**Figure 3:**
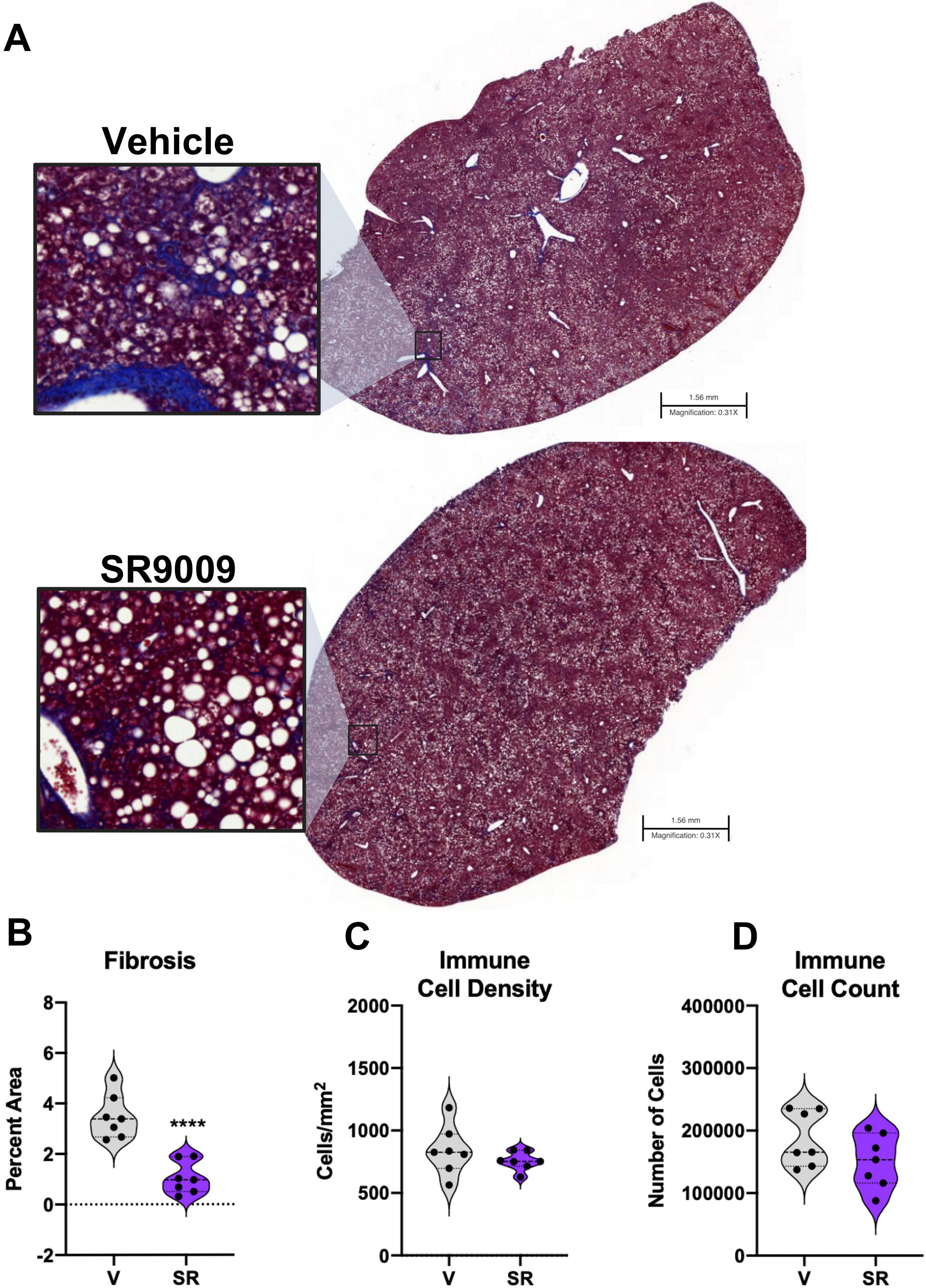
SR9009 Significantly Reduced Fibrosis and Suppressed Inflammatory Activity. (A) Masson’s Trichrome staining of liver sections (representative samples). Vehicle pictured at the top shows significant collagen formation. (B) Quantitated analysis of the Masson’s Trichrome staining for fibrosis as measured by percent of the total area. Immune Cell Density (C) and Immune Cell Count (D) are unchanged, but appear to be less active as fibrosis is significantly reduced in SR9009 samples. Scale bars indicate 1.56 mm.

To continue our investigation into whether pharmacological activation of REV-ERB is beneficial in an NASH model, we analyzed gene expression of inflammatory markers by QPCR from frozen mouse liver tissues. For this analysis, we focused on expression changes related to inflammatory markers that indicate progression of NAFLD towards a NASH pathology, specifically *IL-1*α, *IL-1*β, *Ifn*γ, *and TNF*α. Previous work from our lab [16] and others have shown that REV-ERBs regulate a variety of genes involved in the pathogenesis of metabolic diseases including those associated with inflammation.

Specifically, we were interested to see whether we were suppressing the progression of NAFLD towards NASH with SR9009 by not only alleviating liver damage as assessed in Figure 1D-H, but by also suppressing hepatic inflammation in these animals, which was suggested in the H&E staining (Figure 2). Indeed, when compared to the vehicle-treated group, the SR9009 mice display significantly lower levels of expression of inflammatory cytokines, all of which have been implicated as biomarkers in NASH progression (Figure 4A). Since we observed a significant reduction in fibrosis in the SR9009-treated animals, we also assessed pro-fibrotic gene expression by QPCR. Fig 4B confirms that pharmacological activation of REV-ERB significantly reduces the expression of various pro-fibrotic genes including *Col3A1, Acta2, STAT1, Mmp13, Timp1, and TGF*β [25]. Interestingly, several other pro-inflammatory genes (*Col1A2* and *Agt*) are not affected by SR9009 treatment suggesting that REV-ERB may play a role in the regulation of a select array pro-fibrotic genes involved in NASH progression. While these results indicated a significant effect on pro-inflammatory and pro-fibrotic genes in liver tissues, we sought to assess peripheral inflammation in these animals as well. We assessed plasma TNFα using levels at the time of euthanasia and as shown in Figure 4C, SR9009-treated mice display significantly reduced circulating TNFα levels. This suggests that activation of REV-ERB in a NASH model may dampen the progression of NAFLD towards NASH and improve hepatic health by suppressing fibrosis and inflammatory activity.

**Figure 4:**
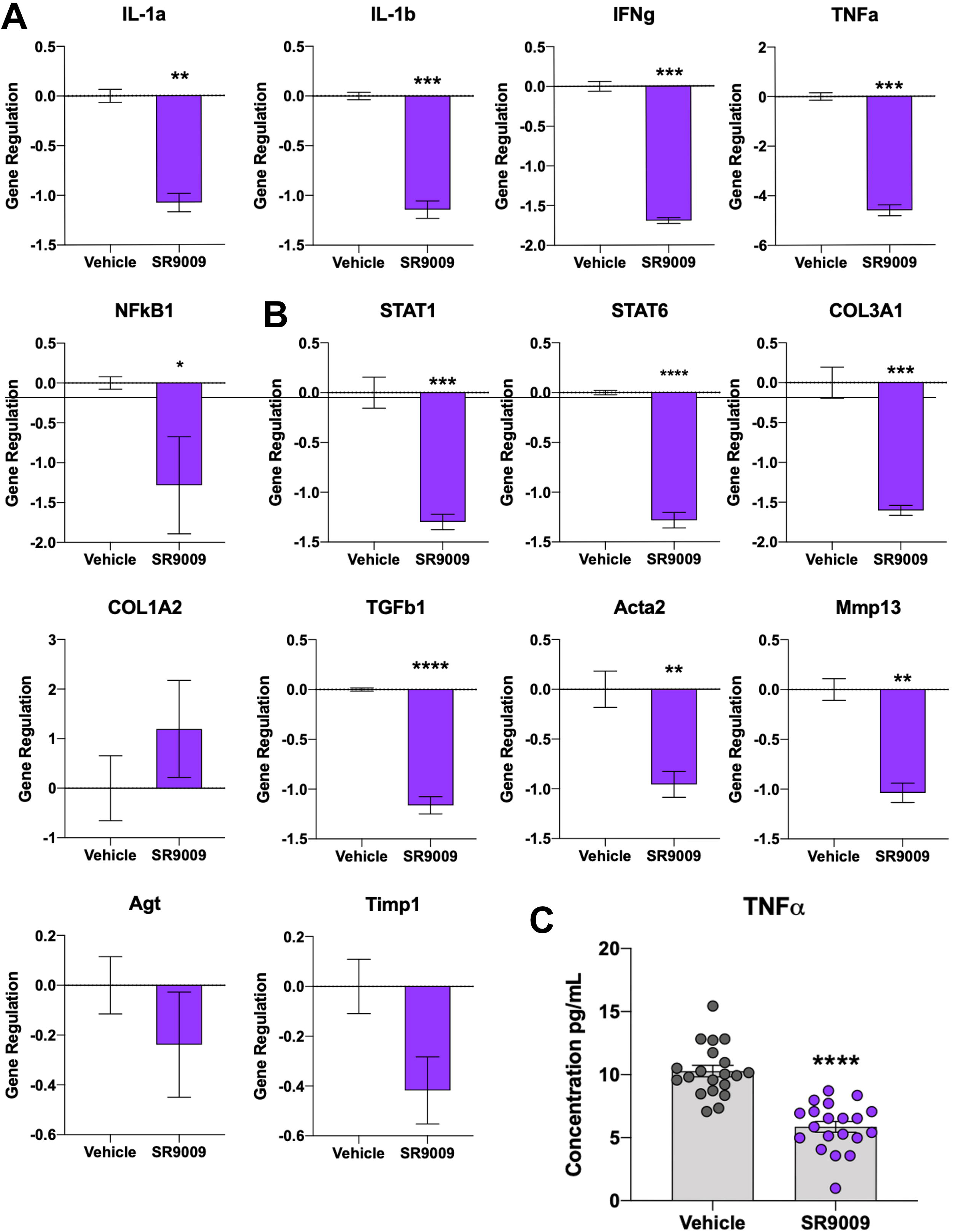
Expression of inflammatory markers are downregulated by SR9009 treatment in a mouse model of NASH. (A) Gene expression of inflammatory cytokines (*IL-1a, IL-1b, IFNg, TNFa*, and *NFkB)* in mouse liver tissues that are involved in the pathogenesis of NAFLD-NASH. (B) Gene expression was also performed for pro-fibrotic genes. As shown in the figure, several genes are significantly downregulated by SR9009 pharmacological activation of REV-ERB. (C) An ELISA for mouse TNFα was performed using plasma samples from each mouse in triplicate and shows that SR9009 mice had significantly reduced circulating TNFα.

## Discussion

Over the last several years, many studies by our lab and others have demonstrated that the REV-ERBs regulate a variety of genes involved in lipogenesis, metabolism, and inflammation (Fig 5). Several studies have investigated potential of REV-ERB agonists as potential therapeutics for cardio-metabolic diseases including atherosclerosis which is a common comorbidity with NAFLD or NASH [16,19]. Understanding the critical role that the REV-ERBs play in these pathways, we hypothesized that pharmacological activation of REV-ERB, with the tool compound SR9009, in a mouse model of NASH would provide beneficial metabolic effects in the liver leading to reduced NASH pathology. While the SR9009 group did not gain weight at the same rate as the vehicle group and we cannot validate whether this was a direct effect of the compound, although feeding and other behaviors were observed to be the same for both groups throughout the study. Our data clearly suggests that SR9009 improved the metabolic profile in the mice (reduced glucose levels and improved glucose tolerance) while also improving hepatic health by suppressing inflammation that essential for the progression of NAFLD to NASH. Most importantly, we observed a significant reduction in expression of pro-fibrotic genes in the liver, which was consistent with reduced hepatic fibrosis.

**Figure 5:**
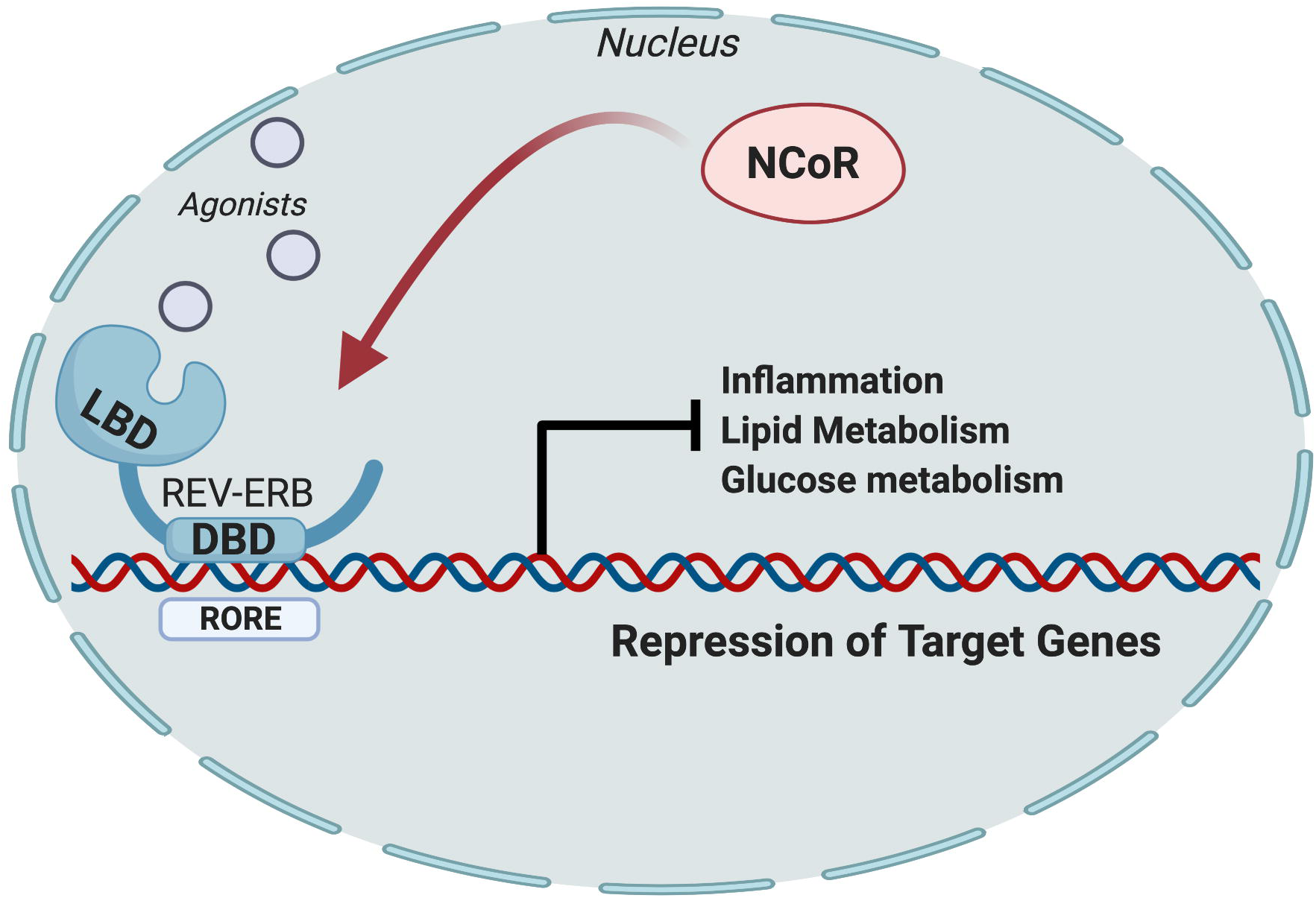
The Nuclear Receptor REV-ERB Regulates Inflammation, Lipid Metabolism, and Glucose Metabolism by Recruiting NCoR to Suppress Transcriptional Activation of Target Genes. This schematic demonstrates how the REV-ERB nuclear receptors regulate transcription of target genes that are involved in inflammation, and lipid and glucose metabolism. Upon agonist binding to REV-ERB, a conformational change occurs and allows for the nuclear receptor co-repressor complex (NCoR) to bind and inhibit the transcription of target genes. As REV-ERBs are regulators of inflammation and hepatic metabolism, it was hypothesized that pharmacological activation of REV-ERB in a NASH model would improve overall hepatic health by suppressing genes involved in lipid metabolism. As indicated by the results, SR9009 actually had little to no effect on lipid metabolism but improved overall clinical indications of NASH in this model. Further investigation demonstrated that REV-ERB agonism in a NASH model suppresses hepatic inflammation and fibrosis and shows therapeutic benefit in dampening the progression of this disease. Image created in Biorender.

Interestingly, we did not observe a significant effect on hepatic steatosis or total lipid accumulation, suggesting that the beneficial effects of REV-ERB agonism in this model may not be due to REV-ERB’s role in reduction of fat mass. Based on the quantitative analysis performed on the liver sections, it appears that hepatocyte ballooning was somewhat decreased in the SR9009 group, although this did not reach significance. This study was performed in a fairly short period of time (30 days) and it is possible that an extended period of dosing may have a more beneficial effect on hepatic steatosis. We originally designed our hypothesis based on our previous study [18] that demonstrated a significant metabolic impact in hepatic lipogenesis and obesity in DIO mice. In this study, SR9009 was dosed twice per day (at 100 mg/kg) and resulted in significant repression of lipogenic and metabolic genes, as well as significant reduction in overall body weight. It is possible that in the current study, the once per day dosing (100 mg/kg) that was utilized to reduce animal stress did not reach the maximal potential efficacy in the NASH model. We utilized SR9009 as a tool to activate REV-ERB in vivo and currently there are limited pharmacological tools available, but in terms of a pharmacological agent SR9009 has relatively poor potency (1 μM range), solubility, and pharmacokinetic properties that lead to a quick clearance time that may attribute to lowered efficacy. Novel REV-ERB compounds with improved potency and pharmacokinetic profiles may provide efficacy in NASH studies to suppress hepatic metabolic activity, inflammation, and fibrosis in the future.

In summary, our data suggests that REV-ERB is a potential therapeutic target to slow the progression of NAFLD towards NASH. The REV-ERB agonist SR9009 displayed efficacy in reduction of hepatic pathology associated with NASH. SR9009 is effective in suppressing clinical markers of liver damage, circulating lipids, hepatic fibrosis and markers of inflammation. Our data suggests that REV-ERB agonists may offer novel therapies for NAFLD or NASH in the future.

## Acknowledgments

We thank Barb Nagel at Saint Louis University School of Medicine (St. Louis, MO) Histology Core for paraffin embedding and performing histological staining of liver sections, Melissa Kazantzis at The Scripps Research Institute in Jupiter, FL for performing clinical chemistry analysis on samples and Sherry Burris for sectioning of liver samples.

## Supporting Information

**S1 Fig. Additional clinical chemistry analysis of plasma from mouse NASH model**. Percentage of Liver weight to total body weight, AST levels, and plasma insulin levels were not significantly changed by SR9009 treatment although total protein was significantly reduced in these mice.

**S2 Fig. REV-ERB target genes involved in cholesterol and lipid metabolism are significantly downregulated by SR9009 treatment in a mouse model of NASH**. We analyzed several REV-ERB target genes (*Dhcr24, ApoeE*, and *ApoC3*) by QPCR to determine expression level differences in the groups. As indicated by the graphs, all three target genes were significantly downregulated, suggesting that SR9009 treatment was suppressing these metabolic pathways but had poor efficacy for reducing steatosis.

**S3 Fig. Whole liver section images showing Masson’s Trichrome staining for fibrosis**. Scale bars indicate 1.56 mm.

**S1 Table. Summary of Histo-Pathological Analysis on Liver Sections**.

## Notes

### Competing Interest Statement

The authors have declared no competing interest.

