## Supplementary figures and images for "REV-ERB Agonism Improves Liver Pathology in a Mouse Model of NASH"

### S1 Fig

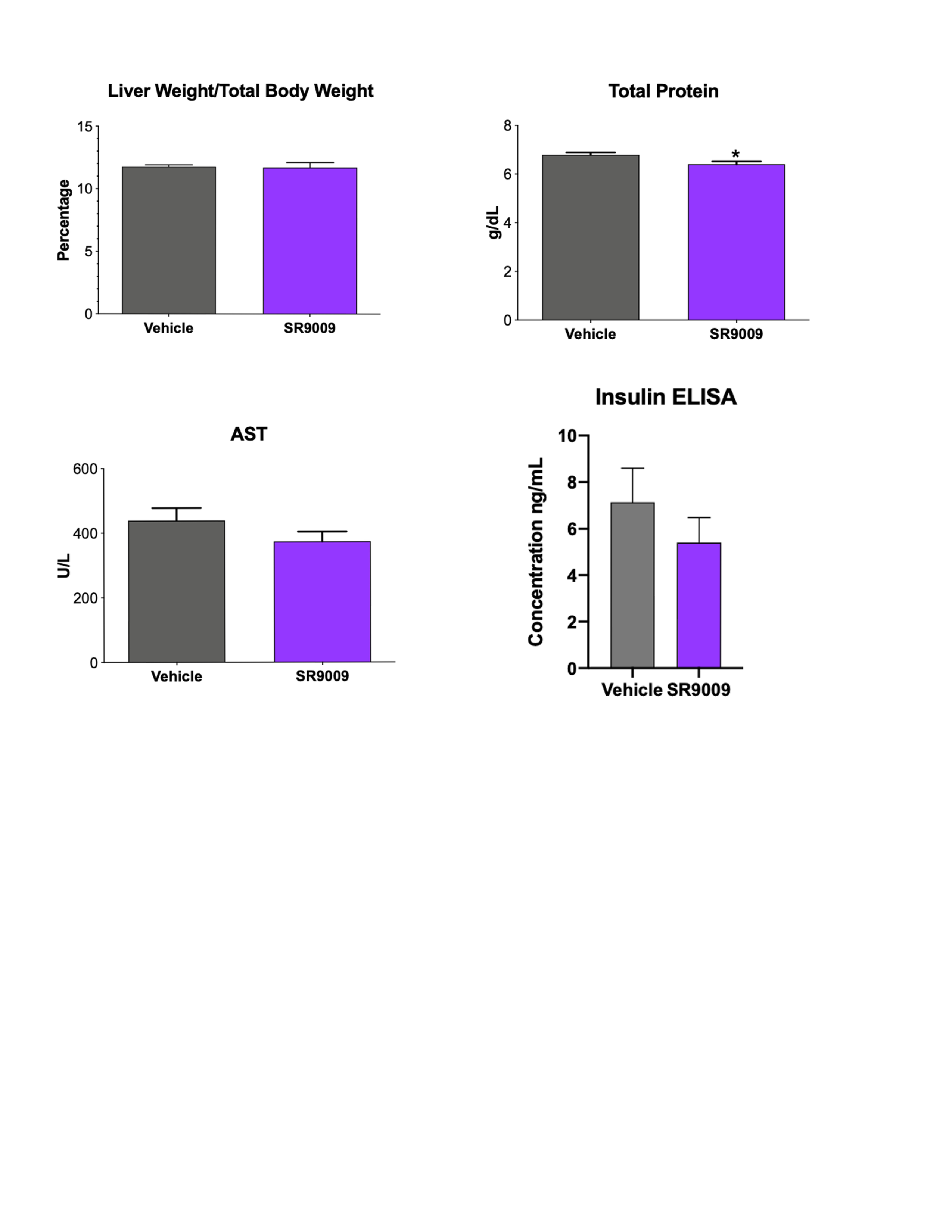

### S2 Fig

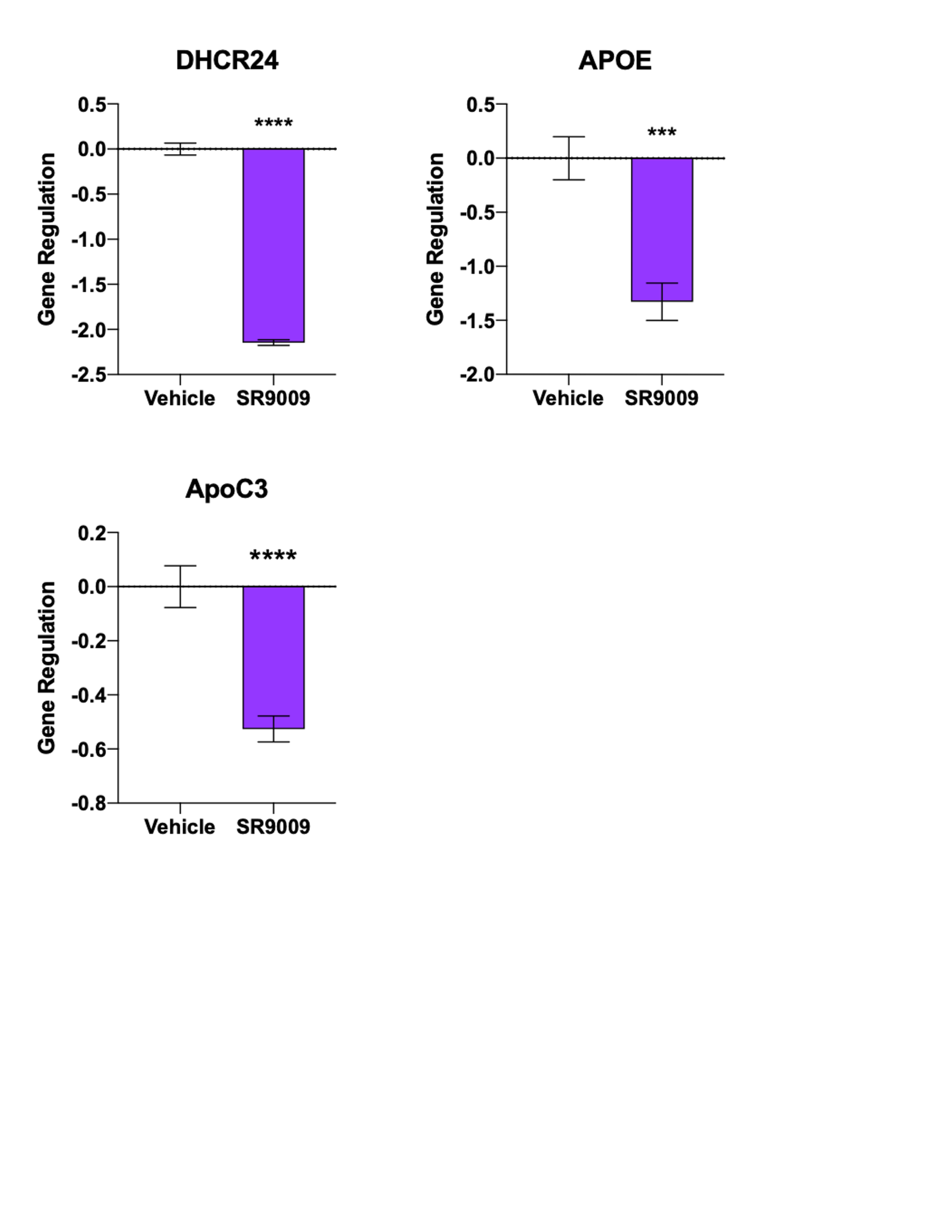

### S3 Fig

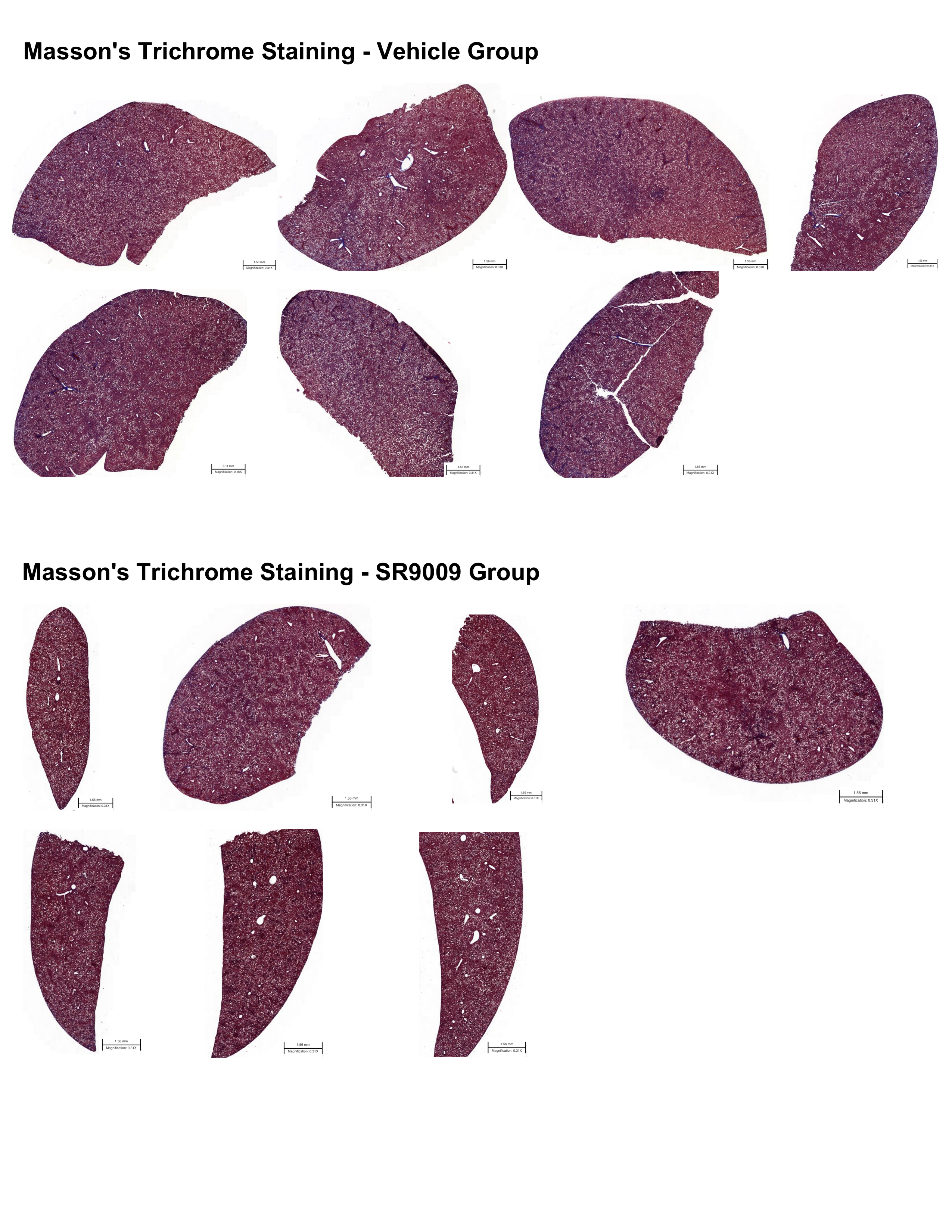
