## Supplementary material for "REV-ERB Agonism Improves Liver Pathology in a Mouse Model of NASH": S1 Table

| **Supplemental Table 1: Summary of Histo-Pathological Analysis on Liver Sections.** | | | | | | |
| --- | --- | --- | --- | --- | --- | --- |
|  |  |  | **Vehicle-Treated** | | **SR9009-Treated** | |
|  | **Steatosis** | | Mean | SEM | Mean | SEM |
|  |  | % Lipid Area | 10.6699 | 0.5175 | 11.1016 | 0.7272 |
|  |  | Total Lipid Area (mm^2^) | 23.76 | 1.3301 | 21.78 | 2.04 |
|  |  | % Macrovesicular | 92.22 | 0.4001 | 92.43 | 0.3693 |
|  |  | % Microvesicular | 7.776 | 0.4004 | 7.664 | 0.3693 |
|  |  | Average Vesicle Size (μm) | 150.9 | 6.649 | 150.9 | 6.573 |
|  | **Ballooning and Mallory Bodies** | |  |  |  |  |
|  |  | Ballooning Hepatocyte Density (Cells/mm^2^) | 78.38 | 5.6537 | 66.84 | 6.3194 |
|  |  | Mallory Bodies Present | Present | | Present | |
|  | **Inflammation** | |  |  |  |  |
|  |  | Immune Cell Density (Cells/mm^2^) | 841.092 | 74.4559 | 755.943 | 28.3342 |
|  |  | Immune Cell Count | 186998 | 16648 | 151180 | 16227 |
|  |  | Total Immune Cell Area (mm^2^) | 20.59 | 1.0573 | 19.05 | 1.9442 |
|  | **Fibrosis** | |  |  |  |  |
|  |  | % Fibrosis Area | 3.479 | 0.3304 | 1.039 | 0.2401 |
|  |  | Total Fibrosis Area (mm^2^) | 7.792 | 0.8168 | 1.9015 | 0.37101 |
